# The local topology of dynamical network models for biology

**DOI:** 10.1101/2023.01.31.526544

**Authors:** Enrico Borriello

**Affiliations:** School of Complex Adaptive Systems, Arizona State University, Tempe, AZ, USA

**Keywords:** Networks, Bioinformatics, Machine Learning, Data Science

## Abstract

The search for motifs –recurrent patterns in the topology of a network– has allowed the identification of universal classes of complex systems from very diverse fields, and has been used as a quantitative tool to highlight unifying properties of evolved and designed networks. This work explores if, and to what extent, network superfamilies previously identified through census of triadic motifs, are represented in the largest data set of dynamic, biological network models. This work presents triad significance profiles of 71 existing biological, and experimentally inspired, network models. The data generated is treated agnostically, and consistently clustered to two classes using several unsupervised techniques. The more populated class correlates with the previously identified superfamily of sensory transmission networks, characterized by the feedforward loop motif typical of signal-processing systems. The other class, surprisingly, better correlates to the superfamily of word-adjacency network. The result is analyzed for varying network size thresholds, and connected to the effect of the model building activity. It is shown that the topology of biological subnetworks starts resembling the topology of “sentences” in word-adjacency networks when the model focuses on smaller portions of the network, coarse-graining the boundary dynamics.

## Introduction

Over the last several decades the scientific community has shown an ever-increasing interest in network theory [1], also thanks to its potential to exhibit shared properties of otherwise vastly different complex systems. Genetic regulation, social interactions, the internet, general principles of linguistics, are just some examples of systems susceptible of a network description, and where this abstraction has proved itself useful [2]. The recognition of the similarities in the network topology of systems as diverse as gene regulatory networks and the World Wide Web is nothing short of spectacular [3]. And, together with similar examples, has paved the way for the understanding of the common mechanisms responsible for the growth and the evolution of such systems [4]. Translating the interactions among the parts of systems into a topological problem, network theory plays a role in complexity studies similar to the one physics plays in understanding natural phenomena. It provides a common language for quantitative studies in fields as diverse as biology, sociology, economics, etc. The best-known example in this sense is the already referenced explanation of the scale-free nature of many natural and evolved networks in terms of the evolutionary mechanism that shapes, together with their growth, also their topology [4].

While the starting point of Barabasi’s and Albert’s theory is a global property of the network topology, i.e. the scale-free nature of the degree *distribution*, Milo *et al*. have focused on the identification of the predominant *local* interaction patterns responsible for fundamental interactions and functions. Relevant to the work presented here is the identification in [5] of the network motifs associated to the directional transfer of information in biological regulatory networks, as opposed to the mutual information processing typical instead of cliques of socially interacting individuals. These two, and two more recurring arrangements of recurrent, co-occurring motifs has led to the identification in [6] of four *superfamilies* of evolved and designed networks (see Methods).

The current study explores if and how well Milo’s superfamilies are represented in *Cell Collective*, the largest data base of biological, experimentally inspired network models [7]. Cell Collectives includes, among others, models of biological cellular processes, including cycles, differentiation, plasticity, migration, and apoptosis. Biological processes in both humans and other animals, as well as viruses, bacteria and plants all have dynamical models counterparts in the data base. Previous analyses of these networks have already allowed:

- A systematic confirmation, on a phenomenological basis, of the theoretical conjecture that living systems exhibit near criticality. 67 of the ever expanding number of networks in the data base are analyzed in [8], and all of them are found to be near critical.
- A formal justification for the surprisingly small number of transcription factors whose overexpression is sufficient to induce mammalian cell change in cellular reprogramming experiments [9, 10, 11, 12, 13, 14, 15]. The dynamics of 49 of the networks in the data base is examined in [16]. All networks are shown to be *easy* to control, and a theoretical justification is provided.

These analyses highlight the main difference between the dynamical models considered in this study, and the topologies considered in [6], where even the smaller networks, like the transcriptional network of *Escherichia coli* [17] with 424 nodes (operons), is larger than the largest model in [7]. This difference deserves a clarification, as it represent the interpretative lens for the findings in this work. While the topology of a network can be studied easily even for networks as large as the WWW –with 5 billion nodes– studying the dynamics of the interaction among these nodes is a problem whose complexity grows exponentially with the number of nodes. Even the simplest gene regulatory network models –where the expression level of each gene is reduced to only two possible values, either expressed or not– implies the existence of 2^*n*^ possible dynamical states for the network, with *n* being the number of nodes. The study of the dynamics is then translated into a topological problem, i.e. finding the attractor cycles (and fixed-points, in particular) of the directed graph representing the state transitions. In this light, a Boolean, dynamical network with just 30 nodes generates a transition graph whose size is comparable to the <u>WWW.</u> This is the reason for the reduced size of the models in [7]. These models often represent subgraphs of much larger network, where the modeling effort is focused on capturing the feedback interactions of the core nodes responsible for the functions and /or expression of cell types the model is focusing on. On the contrary, the peripheral regions of the network and the environmental conditions are often coarse-grained into *input* nodes [16]. This does not just explain the reduced size range for the networks analyzed here (listed in table 1), but also raises the questions of how many of the local topological features of the parent network these sub-networks are retaining. We will address this question analyzing the triad significance profiles of all networks in the data base, and compare them with the expectations set by *Milo’s et al*.*’s* superfamilies.

**Table 1:** List of the network analyzed with their respective sizes.

| ID | size | name |
| --- | --- | --- |
| 0 | 41 | Apoptosis Network |
| 1 | 14 | Arabidopsis Thaliana Cell Cycle |
| 2 | 53 | B Bronchiseptica And T Retortaeformis Coinfection |
| 3 | 22 | B Cell Differentiation |
| 4 | 25 | BT474 Breast Cell Line Long-term ErbB Network |
| 5 | 16 | BT474 Breast Cell Line Short-term ErbB Network |
| 6 | 17 | Body Segmentation In Drosophila 2013 |
| 7 | 33 | Bordetella Bronchiseptica |
| 8 | 67 | Bortezomib Responses In U266 Human Myeloma Cells |
| 9 | 18 | Budding Yeast Cell Cycle 2009 |
| 10 | 20 | Budding Yeast Cell Cycle |
| 11 | 188 | CD4 T Cell Signaling |
| 12 | 18 | CD4+ T Cell Differentiation And Plasticity |
| 13 | 15 | Cardiac Development |
| 14 | 9 | Cell Cycle Transcription By Coupled CDK ... |
| 15 | 34 | Cholesterol Regulatory Pathway |
| 16 | 70 | Colitis-associated Colon Cancer |
| 17 | 5 | Cortical Area Development |
| 18 | 28 | Death Receptor Signaling |
| 19 | 50 | Differentiation Of T Lymphocytes |
| 20 | 104 | Egfr & Erbb Signaling |
| 21 | 28 | FA BRCA Pathway |
| 22 | 23 | FGF Pathway Of Drosophila Signalling Pathways |
| 23 | 15 | Fanconi Anemia And Checkpoint Recovery |
| 24 | 96 | Boolean Network Model for Cancer Pathways |
| 25 | 73 | Glucose Repression Signaling 2009 |
| 26 | 44 | Guard Cell Abscissic Acid Signaling |
| 27 | 25 | HCC1954 Breast Cell Line Long-term ErbB Network |
| 28 | 16 | HCC1954 Breast Cell Line Short-term ErbB Network |
| 29 | 68 | HGF Signaling In Keratinocytes |
| 30 | 24 | HH Pathway Of Drosophila Signaling Pathways |
| 31 | 138 | HIV-1 Interactions With T-Cell Signalling Pathway |
| 32 | 19 | Human Gonadal Sex Determination |
| 33 | 91 | IGVH Mutations In Chronic Lymphocytic Leukemia |
| 34 | 118 | IL-1 Signaling |
| 35 | 86 | Il-6 Signalling |
| 36 | 131 | Influenza A Virus Replication Cycle |
| 37 | 22 | Iron Acquisition And Oxidative Stress Response ... |
| 38 | 13 | Lac Operon |
| 39 | 33 | Lymphoid And Myeloid Cell Specification And ... |
| 40 | 81 | Lymphopoiesis Regulatory Network |
| 41 | 53 | MAPK Cancer Cell Fate Network |
| 42 | 10 | Mammalian Cell Cycle 2006 |
| 43 | 20 | Mammalian Cell Cycle |
| 44 | 12 | Metabolic Interactions In The Gut Microbiome |
| 45 | 16 | Neurotransmitter Signaling Pathway |
| 46 | 19 | Oxidative Stress Pathway |
| 47 | 62 | Pc12 Cell Differentiation |
| 48 | 15 | Predicting Variabilities In Cardiac Gene |
| 49 | 26 | Pro-inflammatory Tumor Microenvironment ... |
| 50 | 24 | Processing Of Spz Network From The Drosophila Signaling ... |
| 51 | 13 | Regulation Of The L-arabinose Operon Of Escherichia Coli |
| 52 | 25 | SKBR3 Breast Cell Line Long-term ErbB Network |
| 53 | 16 | SKBR3 Breast Cell Line Short-term ErbB Network |
| 54 | 31 | Septation Initiation Network |
| 55 | 139 | Signal Transduction In Fibroblasts |
| 56 | 321 | Signaling In Macrophage Activation |
| 57 | 23 | T Cell Differentiation |
| 58 | 99 | T Cell Receptor Signaling |
| 59 | 40 | T-Cell Signaling 2006 |
| 60 | 61 | T-LGL Survival Network 2008 |
| 61 | 18 | T-LGL Survival Network 2011 Reduced Network |
| 62 | 60 | T-LGL Survival Network 2011 |
| 63 | 24 | TOL Regulatory Network |
| 64 | 11 | Toll Pathway Of Drosophila Signaling Pathway |
| 65 | 42 | Treatment Of Castration-Resistant Prostate Cancer |
| 66 | 26 | Trichostrongylus Retortaeformis |
| 67 | 32 | Tumour Cell Invasion And Migration |
| 68 | 18 | Vegf Pathway Of Drosophila Signaling Pathway |
| 69 | 26 | Wg Pathway Of Drosophila Signalling Pathways |
| 70 | 73 | Yeast Apoptosis |

In the next section we summarize the triadic analysis for the unfamiliar reader. We then perform triadic census for each network in the data base, and show that *Milo’s et al.’s* superfamilies are poorly represented. We then check whether the model building artifacts become less relevant once we restrict the analysis to the larger networks. Once the relative number of connections introduced by the model builder starts to become less relevant than the number of connections directly linked to empirical evidence, we start recovering superfamily 1, typical of directional information processing in biology. The networks in the remaining class share instead some of the defining aspects of the topology of sentences from superfamily 4, typical of word-adjacency networks.

## Methods

Network motifs/anti-motifs are local structures that appear unusually often/rarely in a network. Their likelihood is quantified based on their average occurrence in randomizations of the network that preserve the degree of each node. Slight differences are present in the literature about the tresholds and the randomizations involved in the quantitative definition of a motif. This work adopts the definitions presented in [5, 6], and only considers fully connected triads, i.e. fully connected subsets of three nodes. In directed networks, connected triads can take 13 possible topologies, listed and labeled in figure 1 for *i* = 1, …, 13.

**Figure 1:**
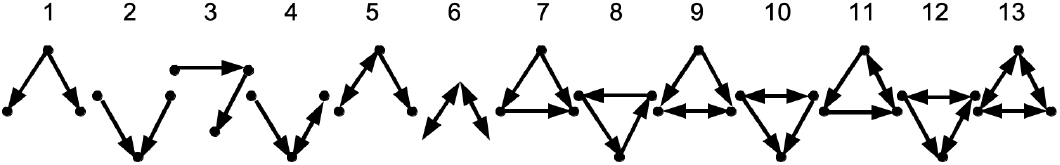
Topologies of the thirteen fully connected triads.

When comparing networks of different sizes, as in this work, a better way to analyze triad censuses is through their significance profiles [6]. For a given network, the occurrence number of triad *i* is denoted by *n*_*i*_, and evaluated in this work using Moody’s matrix expressions [18]. Each count *n*_*i*_ is then compared to the average count 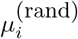 over an ensemble of random networks with the same size and local connectivity for each node. The difference between *n*_*i*_ and 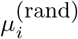 is then compared to 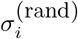 –the standard deviation of 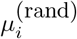 over the random ensemble– and the statistical relevance of the occurrence of triad *i* is parameterized by its *Z*_*i*_ score, defined as

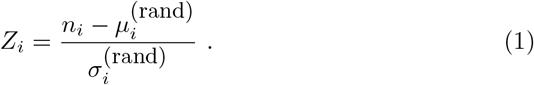

The triad significance profile is then defined as the set of components of the normalized *Z* vector, i.e. SP = *Z/*|| *Z* || _2_, where *Z* = (*Z*_1_, …, *Z*_13_).

Through the analysis of the topology of networks from a wide range of disciplines, Milo *et al*. have identified recurrent significance profiles that they have dubbed *superfamilies* [6]. In this work they will be referred to as superfamily 1 through 4:

**Superfamily 1** It collects sensory transcription networks for the control of gene expression [5, 17, 19, 20]. In these networks nodes represent genes or operons, while the edges describe direct transcriptional regulation. This family is characterized by the positive expression of triad 7, the “feedforward loop” in charge of performing signal-processing tasks [17, 21], and by negative SP values for triad 3, the “3-chain” characteristic of deep networks. This arrangement characterizes the topology of shallow networks capable of fast transient signals [22].

**Superfamily 2** This family also collects biological networks (e.g [23]), but of developmental nature (a smaller subset in *Cell Collective*). The topology of this networks is characterized by positive SP values for triads 7, 9, and 10, and negative SP values for triads 1, 2, 4, and 5. These systems also describe information processing, but with a much slower transient time [24] of the order of hours (as in cell-division) as opposed to mere minutes as in superfamily 1.

**Superfamily 3** It collects networks from the WWW and the internet [4, 25, 26] and it is characterized by positive SP values for triads 9, 10, 12, and 13, and negative values for triads 4, 5, and 6.

**Superfamily 4** This superfamily collects word-adjacency networks from four different languages [27]. Words are parts of speech. Therefore they play specific roles in a sentence, which specifies their position in an adjacency network. This translates in under-representation of *closed* triads (7 to 13), with triad 7 being the most significant anti-motif.

In this work, a similar analysis is performed, and significance profiles (SPs) of triads are derived for 71 experimentally motivated biological networks. These networks, together with their respective size are listed in table. More specifically, for each experimental network in the data base, ten thousand randomizations are performed. Each random network is generated by progressively swapping pairs of edges in a way that preserves both the in-degree and the out-degree of each node (algorithm A of [6]). The total number of switches is equal to 100 times the total number of edges in the network, so that each edge is redirected, on average, 100 times. As in [6], whenever a subgraph appears a very small number of times (less than 2), its count is set to zero.

The significance profiles of all networks analyzed are plotted in figure 2 (light blue lines), together with their average (black, continuous line) and a one standard deviation range (dashed, black lines). Some comments are in order. While the ensemble average is consistent with zero, this must not be wrongly interpreted as lack of motifs. While motifs are present –for *each* real network– based on triad occurrence numbers in network randomizations, the ensemble shown in figure 2 is the totality of real networks. What this figure is showing is that, when the entire data base is considered as a whole, the amount of variance is such that no characteristic profile for the entire data base is highlighted. Nonetheless, figure 2 is not just depicting noise, and denser concentrations of profiles suggest the possibility that a more informative way of plotting these results can be achieved after checking for suitable clusters in the data and splitting the curves accordingly.

**Figure 2:**
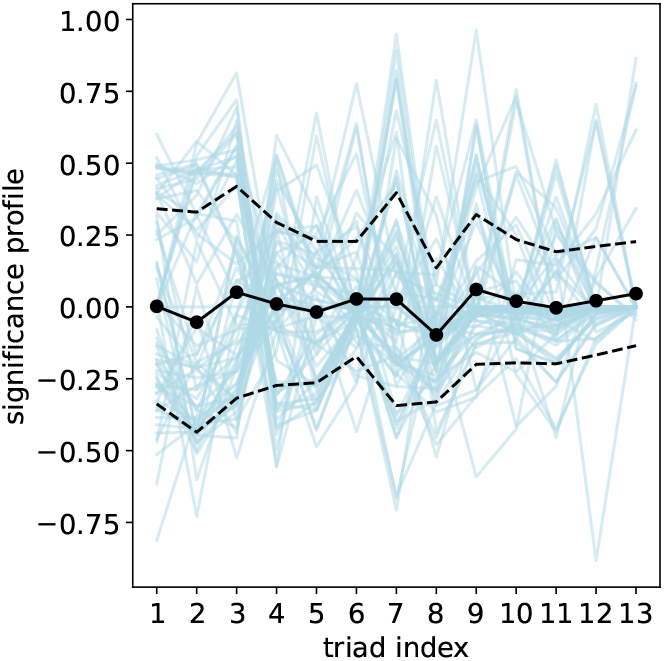
Significance profiles of all networks analyzed. The dashed lines correspond to a 1 *σ* range around the average (black line).

In the rest of this section, several unsupervised clustering and dimensionality reduction techniques are employed to explore whether and how well the profiles from figure 2 can be split into classes, and whether these classes correlate with Milo’s superfamilies.

First, a classification of the SPs is performed in an agnostic way using a K-Means clustering algorithm (more specifically, accelerated K-Means++ [28, 29, 30]) within the 13-dimensional space where each axis corresponds to the normalized *Z*_*i*_ score of triad *i*. In order to select a statically significant number of classes *k*, silhouette scores [31] are plotted in figure 3 (left) for values of *k* from 2 to 10. The silhouette score is the mean silhouette coefficient over the ensemble of real networks, with each coefficient measuring how well each SP is labeled based on its Euclidean distance from its cluster and the distance from the nearest foreign cluster. Each coefficient, and therefore the score, can vary from *−*1 to +1, with +1 corresponding to perfectly separated, and far away clusters. Given the amount of variance present in figure 2, it is not surprising that the largest score is as low as *∼*0.34. Nonetheless, this value is significantly greater than for any other value of *k*, and corresponds to a split of the data base in two clusters

**Figure 3:**
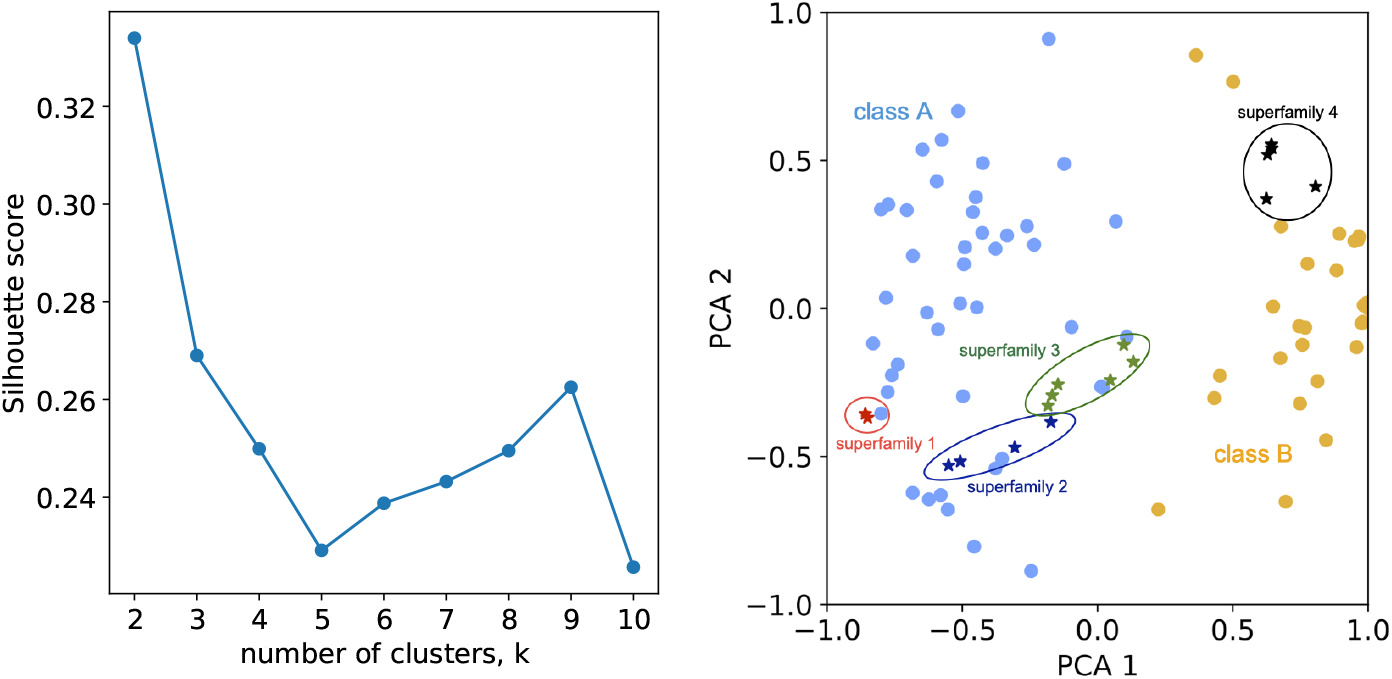
(Left) silhouette scores for varying numbers of clusters *k*. The data set is best split in two clusters: class A and class B. (Right) PCA of the 2 clusters. Blue dots correspondd to class A, yellow dots to class B. The stars correspod to projections of the significance profiles of the networks analyzed in [6]. Notice that networks belonging to the same superfamily are clustered within the same class.

To visualize the two classes, the SPs are projected onto the 2-dimensional space spanned by the first two coordinates of a principal component analysis. Different colors in figure 3 (right) correspond to different classes, and visually confirm the outcome of the K-Means analysis. This visualization also offers another advantage, as it allows the projection of the SPs of the networks analyzed in [6] onto the same plane. Interestingly, Milo *et al*.’s SPs belonging to the same superfamily are not scattered *across* classes. But, before discussing the meaning of this, one last 2-dimensional visualization and clustering technique is proposed, in the form of a heat map of the correlation matrix (as in [6]). Once the ordering of the networks is rearranged to maximise correlation within the diagonal blocks, two classes emerge in figure 4. The two classifications are found to be in good agreement with one another.

**Figure 4:**
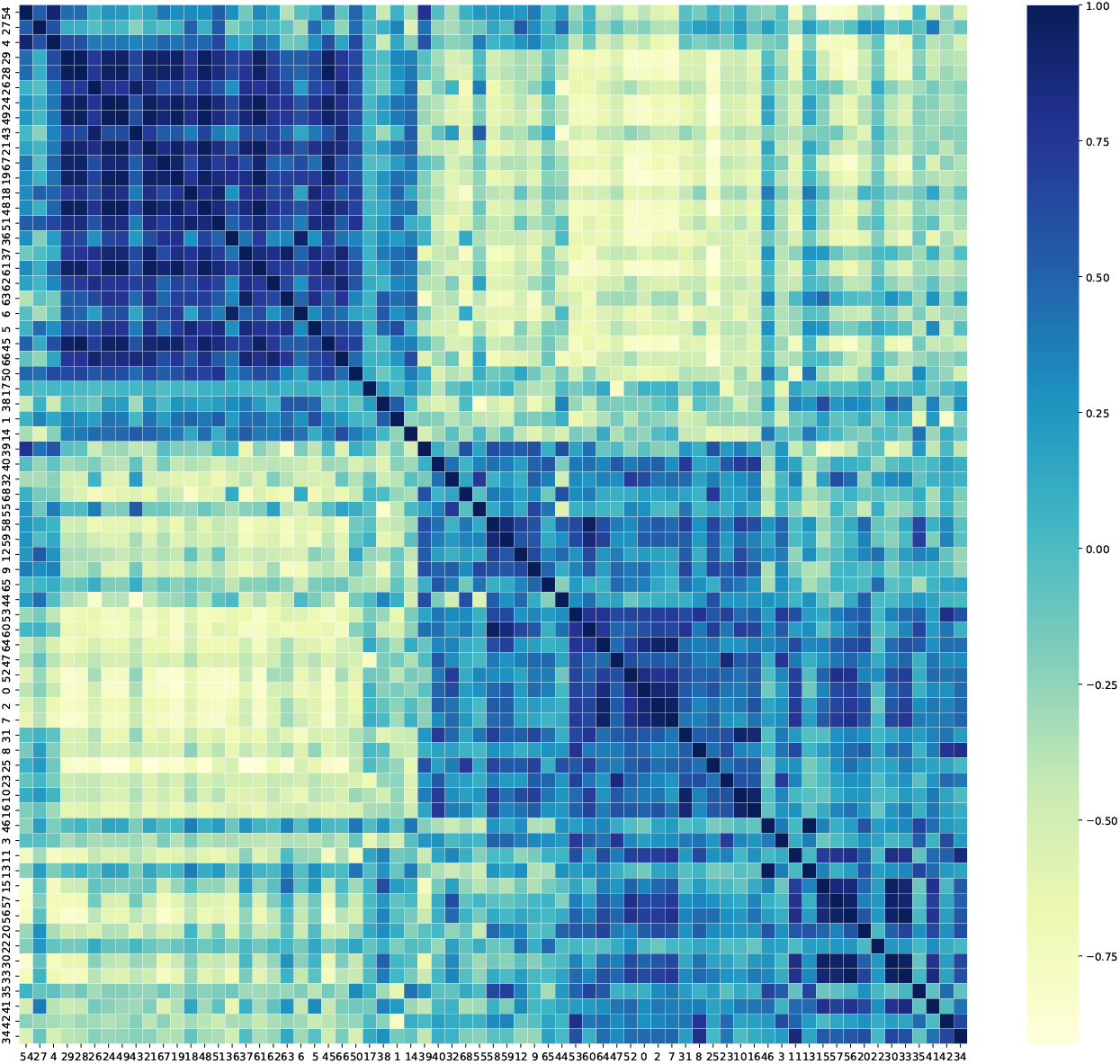
Correlation coefficient matrix of the triad significance profiles for all the dynamical networks analyzed.

Now that more confidence in the distinct nature of these two classes has been established, their differences can be better understood and analyzed. Figure 5 splits the content of figure 2 in two panels, each one showing the SPs of just one class. More recognizable profiles start to emerge, with vertical black lines highlighting regions no longer consistent –at a 1 *σ* confidence level– with their random counterparts. As expected, the first feature this split is highlighting is the know conservation law which anti-correlates the occurrence number of triangular and linear topologies, i.e triads where all three and only two of the three nodes are directly connected [6].

**Figure 5:**
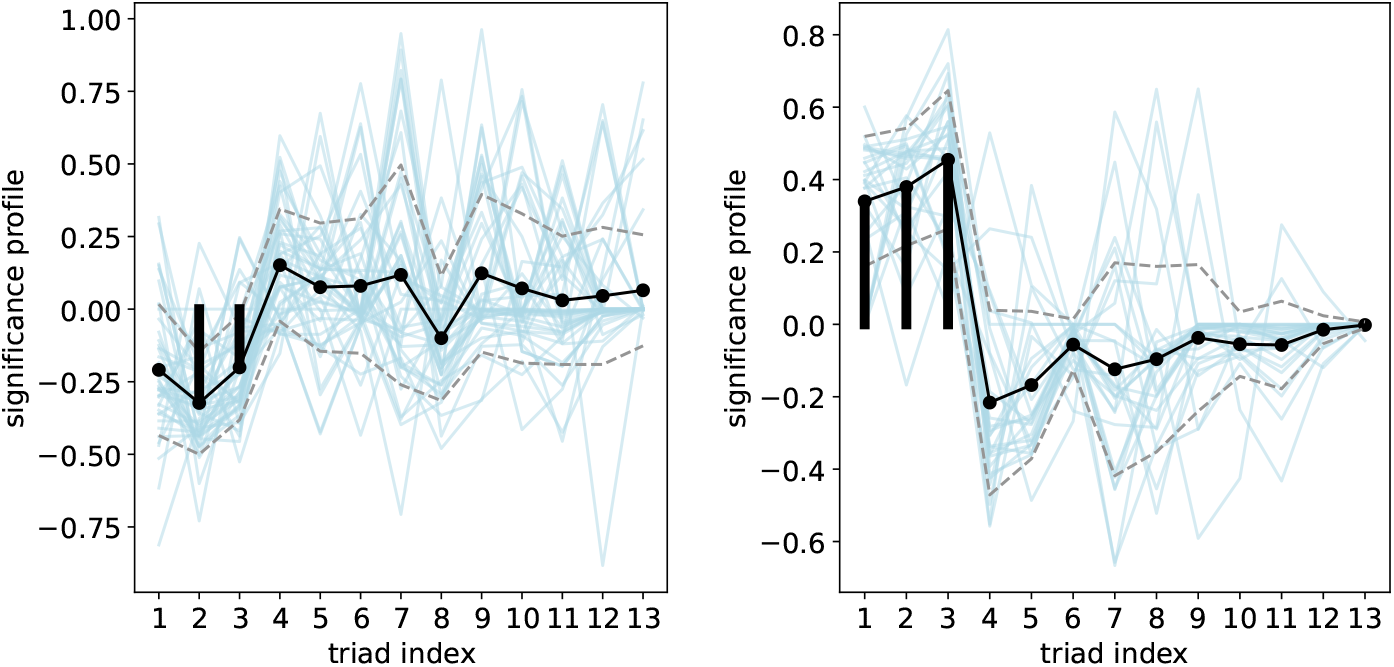
Significance profiles of all networks analyzed. Profiles corresponding to different classes are plotted separately (A to the left, B to the right). The dashed lines correspond to a 1 *σ* range around their respective average (black lines).

Consistently with what found in figure 3 (right), the less populated of the two classes (27 out of 69 networks, right panel]) begins to show a surprising similarity with *Milo’s et al*.*’s* superfamily 4, typical of word adjacency networks. The more abundant class (remaining 42 networks, left panel) seems to correlate better with the first three superfamilies (again, in agreement with figure 3 (right)). Nonetheless, figure 5 fails to exhibit the discriminant role played by triad 7 — the feedforward loop typical of signal-processing systems– and an anti-motif of word-adjacency networks. The more populated class in figure 5 will be denoted class A, while the remaining class will be referred to as class B.

In what follows we want to explore the conjecture that a better representation of superfamily 1 (the expectation for the data analyzed) can be recovered after the effect of of the model building activity on the topology of the network is reduced. The main assumption is that this effect is greater in smaller network models. As the model focuses on progressively smaller portion of the biological network, larger and peripheral regions of the system are coarse-grained to a reduced number of *input* nodes, constituting the effective boundary conditions, and a subset of the control knobs of the dynamics of the group of nodes at the core of the model [16].

The previous analysis is therefore repeated after setting a threshold on the network size at 25 and 50 nodes respectively. For higher thresholds, the statistics starts to become insufficient, with just seven networks with more than 100 nodes. At that point, the specifics of the selected networks become more relevant than their average, and no longer representative of the original ensemble. Even after restricting the data set to just the large networks, *k* = 2 is still the most significant number of classes, as demonstrated by silhouette scores analogous to the ones in figure 3 (left). The number of networks *correctly classified*, i.e. belonging to class A, increases from 59% (41/69 networks) to 71% (26/40) when only networks with *n ≥* 25 nodes are considered, and to 75% (18/23) for *n ≥* 50. Profiles for *n ≥* 50 are shown in figure 6, where they are also compared to the average profiles of Milo’s superfamilies. The most distinctive difference between figures 6 and 5 is the more prominent role played by triad 7, further corroborating the similarity between class B and Milo’s superfamily 4. Better separated classes are now also visible in figure 7 where the correlation matrix of the SPs, is derived for just the networks with *n ≥* 50.

**Figure 6:**
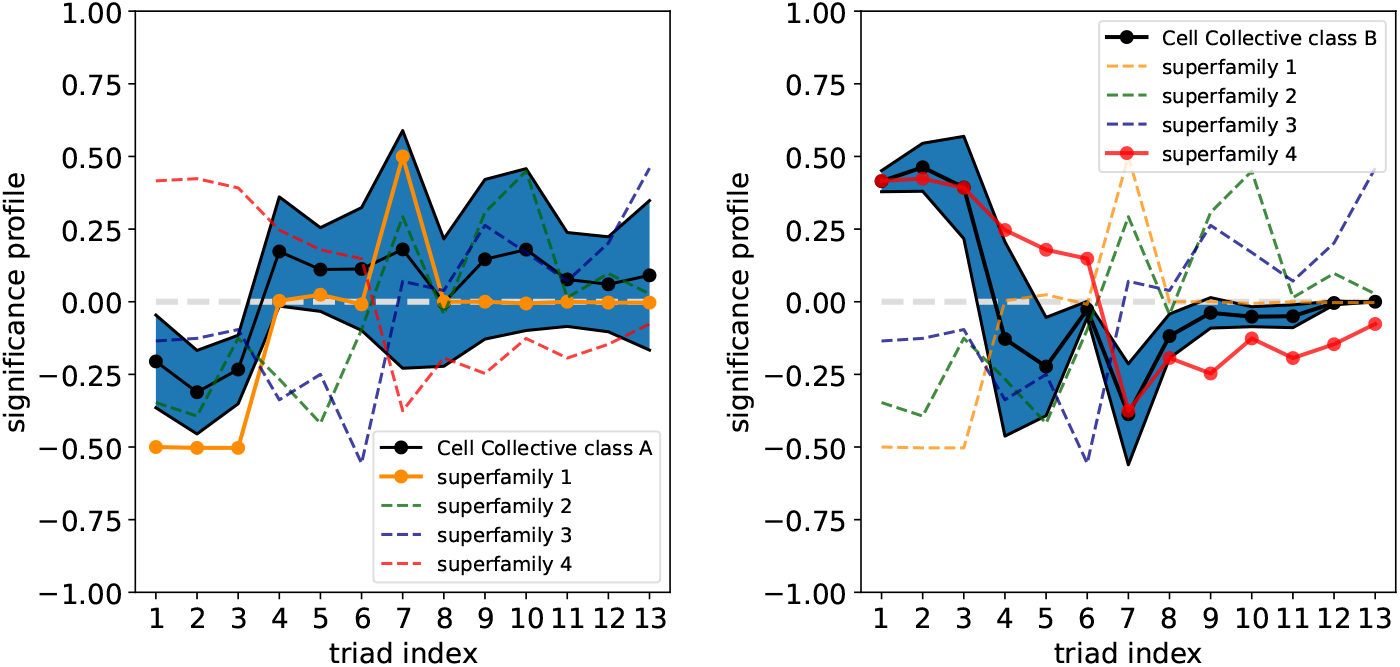
Comparison between the average significance profiles for class A (left) and B (right) with the average profiles of *Milo’s et al*.*’s* superfamilies. This panels only include networks from our data set with size greater than or equal to 50 nodes. Notice the much more polarizing nature of triad 7.

**Figure 7:**
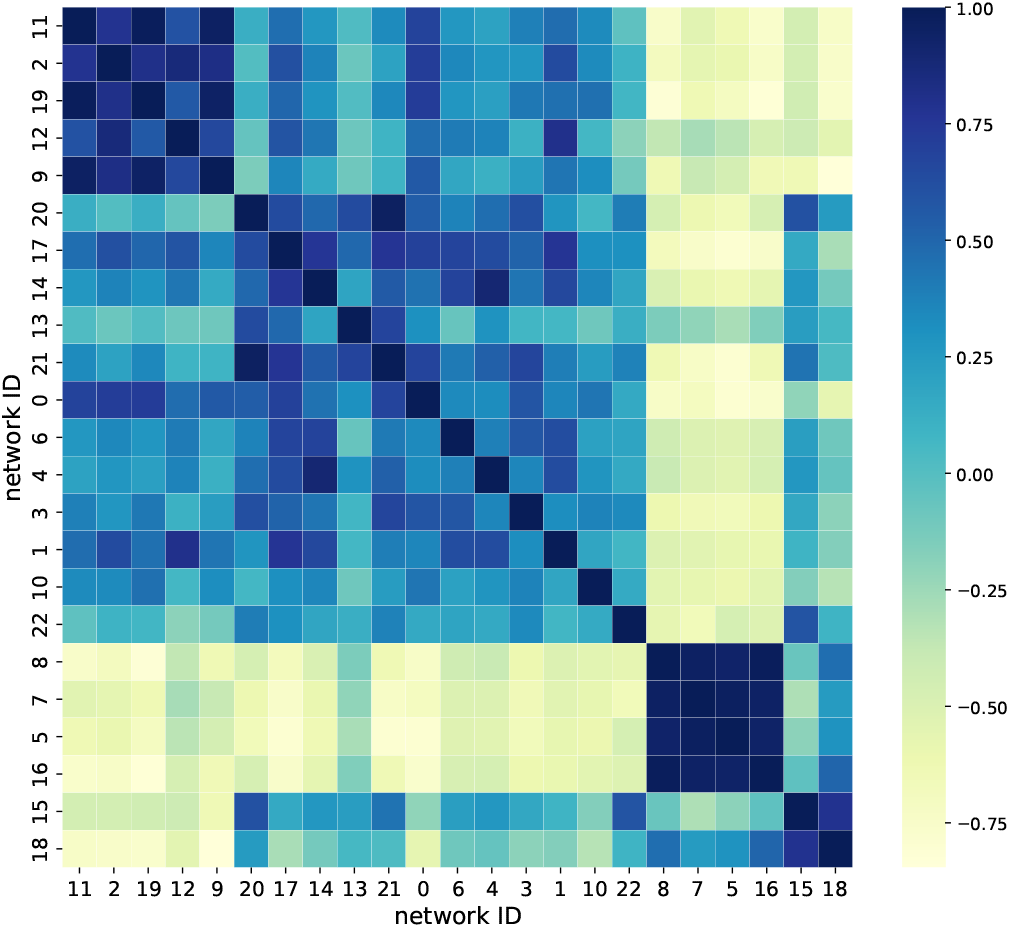
Correlation coefficient matrix of the triad significance profiles for just the networks with size greater than or equal to 50 nodes.

Table 2 collects the correlation coefficients between the average SPs of classes A and B with the averages SPs of Milo superfamilies 1–4 for all networks, only networks with *n ≥* 25 and *n ≥* 50 respectively. Highlighting values greater than 0.5 (boldfaced in the table) is enough to show that –regardless of the threshold on the network size– superfamily 1 consistently correlates to class 1, while superfamily 4 is always well correlated to class B. Supeperfamilies 2 and 3 do not show significant correlation with either class. Notice that, while there is no significant trend in the improvement in the correlation coefficients for increasing values of the threshold, the percentage of networks belonging to class A does improve, as noted before.

**Table 2:** Correlation coefficient between the average significance profiles of class A and B with the average profiles of *Milo’s et al*.*’s* superfamilies for increasing size of the size threshold. Values greater than 0.5 are boldfaced, and consistently show that, regardless of the threshold on the network size, superfamily 1 is always well correlated to class 1, while superfamily 4 is always well correlated to class B. No other significant correlation is found with supeperfamilies 2 and 3.

| | all networks | | $n \geq 25$ | | $n \geq 50$ | |
| --- | --- | --- | --- | --- | --- | --- |
|  | class A | class B | class A | class B | class A | class B |
|  | 62% | 38% | 70% | 30% | 78% | 22% |
| superfamily 1 | **0.84** | -0.87 | **0.92** | -0.91 | **0.88** | -0.96 |
| superfamily 2 | 0.51 | -0.35 | 0.57 | -0.39 | 0.57 | -0.45 |
| superfamily 3 | 0.13 | -0.032 | 0.18 | -0.058 | 0.16 | -0.11 |
| superfamily 4 | -0.63 | **0.65** | -0.74 | **0.72** | -0.71 | **0.75** |

## Discussion

Comparative analyses of the local topology of large designed and evolved networks has allowed the identification of recurrent arrangements of motifs with high explanatory power. The census of triadic motifs constitutes the most prolific of these strategies. It has allowed the identifications of the characteristic triadic profiles that summarize: The logic evolved in biological regulatory networks to process information at adaptive speed (*Milo’s et al*.*’s* superfamilies 1 and 2); the topology of social interactions (superfamily 3); The recurrent patterns in the way the different parts of speech are arranged in several natural languages (superfamily 4) [6]. This work explores whether these previously identified classification is consistent with the largest data set of dynamic, biological network models to date [7].

The expectation of a strong representation of superfamily 1 and –to a less extent– 2 is not met unless the analysis accounts for the choices of the model builder. While the topology of large gene regulatory networks can be inferred through gene knockout techniques [32], only much smaller portions of these networks can be modeled as dynamical systems, because of the exponential growth of the state space of the dynamics with the size of the network. Upon analyzing 71 existing biological, and experimentally inspired, network models from [7] –and deriving their triad significance profile (SP)– the presence of two distinct clusters in the SPs is consistently highlighted. The number of clusters is deduced agnostically and in an unsupervised way.

Once the data set is partitioned this way, the most populated of the two classes, denoted class A, does correlate with superfamily 1 with a Pearson coefficient of 0.85 –as expected– by virtue of the high number of sensory transmission networks in [7]. While this result is encouraging, and shows that meaningful topological aspect of the real biological networks are retained in many of their dynamical model counterparts, the remaining class, denoted class B, is far from being a negligible subset, and accounts for about 41% of the networks analyzed. Interestingly, among the four superfamilies, class B best correlates with superfamily 4, although with a smaller Pearson coefficient of 0.7. The identity of class 4 –typical of word-adjacency networks– further corroborates the hipothesis that the misclassified networks are the ones most affected by the model building assumptions. Sentences have orderly structures, where some parts of speech always precede others. In building a dynamical model, peripheral regions of the network get coarse-grained, and turned into environmental and input nodes. This adds depth to the topology of the dynamical network, and increases the relative count of triads 1 though 3. To further explore this conjecture, the analysis has therefore been repeated for varying thresholds on the network size, under the assumption that the model building activity is more untactful in smaller models. Unsurprisingly, no significant improvement is observed in the correlations of class A with superfamily 1 and of class B with superfamily 4, due to the much larger variance affecting our data set that the reduced number of representative networks in [6]. But the relative number of networks incorrectly classified is greatly reduced from 41% to 25% once the threshold is raised to fifty nodes.

This work is a first attempt at bridging the interpretational gap between purely topological and dynamical network studies. In particular, it highlights misconceptions that rushed interpretations about the topology of individual networks designed for dynamical analyses might induce. While individual dynamical models can only provide partial and sometimes contrasting pieces of information about topology, the statistical analysis presented here of a large data base of such models shows that the key features of the expected topology of information processing systems is indeed inherited, albeit sometimes hidden by the effect of the model building activity, especially when small networks are considered.

